# Affordable *Caenorhabditis elegans* tracking system for classroom use

**DOI:** 10.1101/2020.12.28.424585

**Authors:** Nicholas Leonard, Andrés G. Vidal-Gadea

## Abstract

For decades the nematode *C. elegans* has served as an outstanding research organism owing to its unsurpassed experimental amenability. This advantage has also made this tiny worm an attractive vehicle for science instruction across higher learning institutions. However, the prohibitive cost associated with the automated behavioral assessment of these animals remains an obstacle preventing their full adoption in undergraduate and high school settings. To improve this situation, we developed an inexpensive worm tracking system for use by high school interns and undergraduate students. Over the past two years this tracker has been successfully used by undergraduate students in our introductory Cell and Molecular lab (BSC220) at Illinois State University. Here we describe and demonstrate the use of our inexpensive worm tracking system.

## Description

From the beginning of *Caenorhabditis elegans*’ inception as a genetic model organism by Sydney Brenner (Brenner 1974), the ability to measure and quantify behavior in these nematodes led to numerous and powerful insights (Apfeld *et al.* 2018). The experimental amenability of worms makes them not just superb research subjects but also useful pedagogical tools. While excellent classroom additions to illustrate many biological processes, the educational potential of worms has lagged due to the expensive equipment required for their study. In most educational settings lack of equipment discourages individual exploration and falls short of the promise owed to young and driven students. One of the places where this is felt strongly is in the automated quantification of animal behavior. Many systems have been developed over the years and are currently used across the world for the rapid and unbiased quantification of behavioral phenotypes. For example, we have used tracking systems in the past to compare the ability of mutant strains to transition between gaits (Vidal-Gadea et al., 2011), and Deng and colleagues to study the role of inhibition during locomotion (Deng et al., 2020, see Husson et al. 2012 for a review of the use of tracking systems in C. elegans). Until recently, the automated quantification of *C. elegans* behavior was only feasible for a few wealthy (or specialized) labs. Recent advances have begun to reduce the expense and complexity of automatically quantifying animal behavior. For example, the Haspel lab at NJIT made use of the recently developed Tierpsy behavioral software (Avelino *et al*. 2018) to build numerous animal tracking systems that are at once user friendly, and able to achieve levels of kinematic analysis that previously required considerably more expensive setups (Deng *et al.* 2020).

After building and witnessing the ease of use and power of the setup described above, we decided to reverse engineer a similar worm tracker system for classroom deployment. Our intention was to develop a bare-bones system that allowed as many students as possible to engage in individualized research projects. We implemented this worm tracker in an introductory Cellular and Molecular Laboratory course at Illinois State University (BSC220). In this class, small groups of (three) students built and regularly used the trackers to film and quantify the behavioral phenotypes of *C. elegans* worms illustrating different cellular or genetic manipulations. Because lab fees associated with molecular labs can be restrictive, a key consideration in our approach was cost reduction (along with ease of build and use). Here we describe an affordable (~US$30) worm tracking system whose construction and use is amenable to high school and undergraduate students. We provide a list of materials, step by step assembly instructions, and illustrate its ease of construction and use to measure kinematic differences between wild-type and mutant *C. elegans* worms.

The process of filming and analyzing animal behavior involves three distinct components. First is the construction of a worm tracking system which is the focus of this manuscript. The worm tracking system needs to be able to image nematodes at sufficient magnification and contrast to allow machine vision algorithms to locate and track individual animals. While inexpensive USB microscopes are available for purchase (and indeed are used in this build), one key feature of useful worm trackers is their ability to provide optimal underneath illumination in order to produce homogeneous, high-contrast, images. The second component of the system consists of an image acquisition program used to film the animals. There are several free programs available including micro-manager, Widows Camera, and the programs distributed with the USB microscopes such as the one used in our tracker. The third and final component of the ensemble is a worm tracking program able to detect and track individual nematodes from a movie or series of images. Presently Tierpsy is an outstanding free kinematic package with widespread use among worm labs (Javert et al. 2018). However, because of its restrictive system requirements we opted for a more limited but easier to install and use ImageJ plugin named wrMTrck (Nussbaum-Krammer *et al.* 2015). The wrMTrck plugin is able to extract many useful parameters such as animal size, velocity, and body bend frequency.

## Methods

### Methods

#### Worm Tracker Construction

Refer to Figure 1A for a series of images illustrating the assembly steps described below.

1. Print and glue the stencil (Supplementary Figure 1) to a sheet of corrugated cardboard (thicker cardboard is harder to cut but provides a stronger build).
2. Use an x-acto knife (or scalpel) to cut the pieces to be used in the tracker following directions on stencil.
3. Prefold pieces (where indicated), begin assembly by folding the tracker base (A).
4. Glue the USB light braces (B) to secure the USB light beneath the tracker making sure the light is aligned with the circular hole on the base, and that the LED faces upwards (through the hole). The USB plug from the LED light should protrude through the rectangular hole cut in the back of the base.
5. Use glue on (C) to close the base up.
6. Prefold and glue the scope towers (D) along the longest sides of the base.
7. Glue the tower brace strip (E) to the top of both tower arms to provide structural reinforcement.
8. Cut and glue the worm plate holders (F) onto the tracker base.
9. Cut a piece of diffuser cloth and glue it to the diffuser holder (G) (a kimwipe or tissue paper will serve as a diffuser if fabric is not available).
10. Wrap the Inner microscope holder (H) tightly around the top of the USB scope and glue it onto itself, creating a tight sleeve for the scope. Make sure to leave the USB light dial (above) and the focus dial (below) on the microscope free to rotate.
11. Fold in half each of the outer scope holders (I) gluing the ends to the inner holder.
12. Insert the folded ends of the microscope holders through the cut in the towers; they should be tight but allow vertical adjustment.
13. Connect the USB light to a USB LED dimmer, and this to a USB power outlet. The USB dimmer allows to control the amount of light hitting the setup which is important to obtain homogeneous illumination.
14. Place a plate with nematodes on the plate holder and connect the microscope to a windows PC with preinstalled acquisition software, ImageJ, and the wrMTrck plugin to start using the setup.

**Figure 1.**
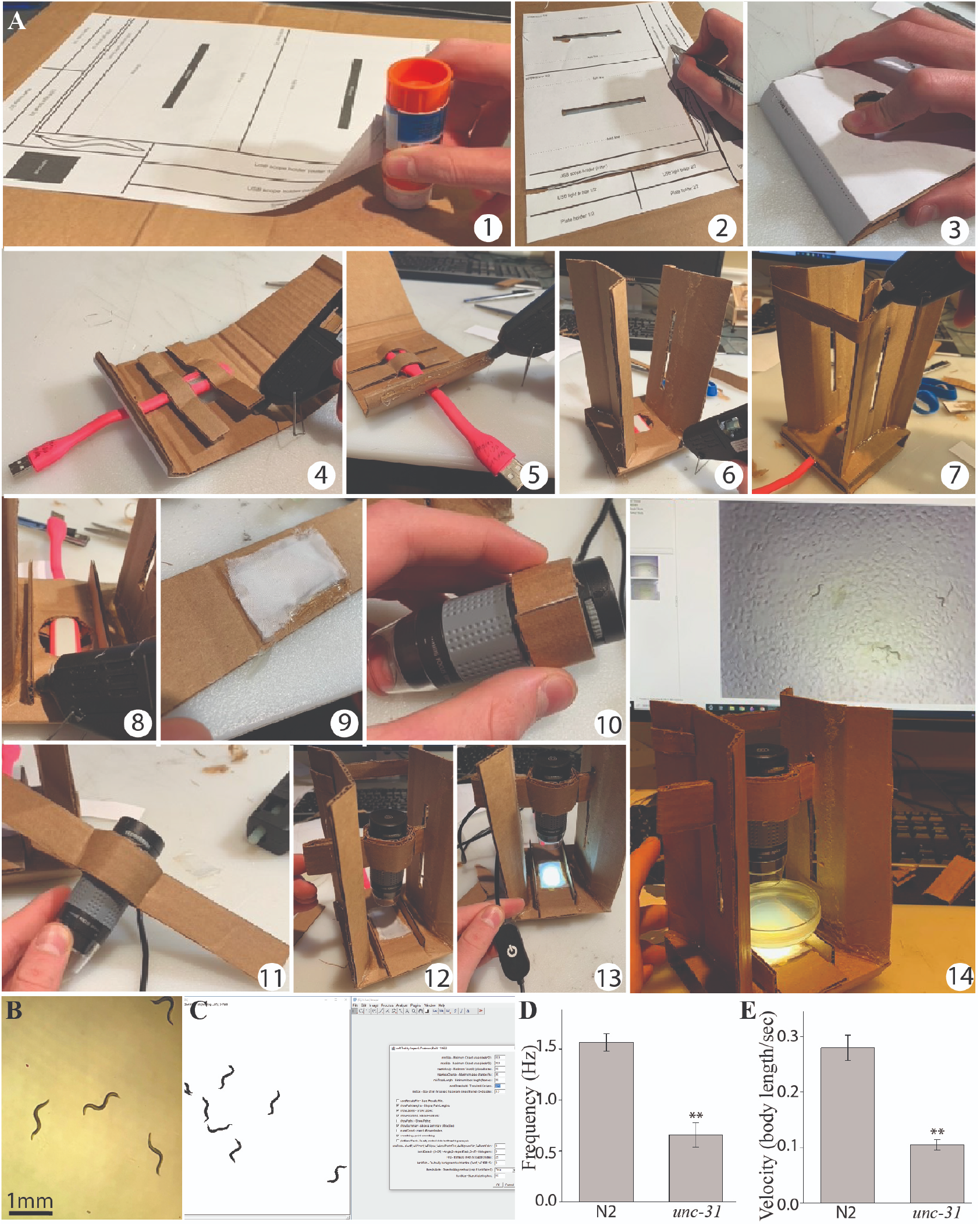
Construction and use of low cost *Caenorhabditis elegans* tracking system. **A)** Assembly of the worm tracker takes under one hour (1-14). See step by step instructions below. **B)** Sample frame from movie obtained with the tracker. **C)** Movie processing with the wrMTrck plugin for ImageJ. We filmed and compared swimming in day 1 adult wild-type (N2) and *unc-31(e928)* mutants to illustrate the capability of the tracking system. Four swimming assays with eight animals each were performed for each strain. **D)** The synaptic mutant *unc-31(e928)* has significant impairments in swimming frequency and normalized (to body length) velocity **(E)** compared to N2 worms. ** p<0.001, t-test.

Depending on the materials used and number of units built, the instructions above will produce worm tracking systems costing as little as US$30 (see list of parts below). We feel that this is a permissive cost in the context of most secondary and tertiary educational settings. Several upgrades can be implemented to increase the longevity (and cost) of the tracker. For example, replacing the cardboard with plastic or wood would produce a much stronger and longer lasting device. For the sake of minimizing difficulty and cost of the build we show here the most inexpensive and simplest configuration. The tracker described here was successfully constructed and used by our BSC220 students and by high school interns in our lab.

#### Image Acquisition

Once the worm tracking system is assembled and connected to a windows machine, the acquisition software provided by the microscope manufacturer can be used to film freely behaving animals. For the example presented here we used *Windows Camera* which comes preinstalled in all Windows PCs. We should mention that the tracking software (ImageJ) is capable of importing a wide array of image and movie formats. However, depending on the software used, the user might need to convert their image or movie files into one of the standard formats used by ImageJ (e.g. TIFF and Jpeg images, AVI movies, etc.). Once the software is started, the user simply needs to select the camera and select the filming option on the control ribbon. The US$20 USB microscope we tested comes with a calibrating ruler (used to establish the magnification), and with an Android phone connector which enables students owning this type of device to record movies without the need for a computer. Before filming, the diffuser slide positioned below the plate of worms. The diffuser ensures that the LED light is evenly distributed throughout the plate. Uniform lighting is of paramount importance during filming. A plate with worms to be filmed is placed above the diffuser. It is best to not attempt filming too many worms at one time, as this increases the likelihood of animals colliding (which complicates tracking). In our experience 6 to 8 worms is ideal. A copper ring can be placed on the plate to keep all the animals within a small area in the field of view (Simoneta and Golombek 2006). We usually film through the lid of the plate but this can also be removed (if reflections appear on its surface). We never use the LED lighting that comes onboard with the USB microscopes. This type of illumination from above is not optimal when filming transparent worms as it results in very poor contrast. The microscope has a red button that allows the user to switch between focus and magnification when turning the silver dial present. The focus dial on the microscope is used to bring animals into focus. The LED light installed underneath the tracker is controlled using the LED dimmer it is connected to. Optimal results are obtained when plates are devoid of any visible imperfections that might challenge the animal detection plugin. To compare worm strains with different locomotor abilities we used eight animals and filmed for 60 second movies at 23fps. Healthy wild-type worms move at about 1cm/min using body bends ranging in frequency between 0.5-2.5 Hz depending on the substrate (Vidal-Gadea *et al.* 2011). Therefore, the parameters used by our setup are appropriate for most rudimentary analyses. When filming, we recommend making multiple movies in anticipation of possible challenges during the analysis step. Movies of worms crawling on agar plates, or swimming in liquid nematode growth media are available in the supplementary section (Supplementary videos 1 and 2).

#### Animal Tracking

Once the animals have been filmed and avi files generated, they can be imported into the analysis software (ImageJ) in order to track the worms and generate descriptive statistics. As we mentioned above there are many available tracking packages, including a few freely available ones. For use in a research lab setting, Tierpsy is likely the best option providing a plethora of useful parameters that include centroid-based metrics (e.g. velocity), and posture-based metrics (e.g. body curvature). An alternative that is more amenable to a high school or undergraduate lab setting is the wrMTrck plugin for ImageJ written by Jesper Pedersen (Nussbaum-Krammer *et al.* 2015). Among the useful parameters measured by this plugin are body lengths travelled per second, body bend frequency, speed, and animal size. These metrics are sufficient to distinguish populations that perform differently due to a plethora of possible experimental manipulations.

#### Evaluation of the tracking system

##### Camera

To evaluate our system’s capabilities we filmed worms under different conditions. The camera manufacturer manual indicates the scope is capable of magnifications between 40x and 1000X, up to 30fps, and 5 megapixels resolution. Magnification can be controlled using a dial on the scope (this also requires changing the height of the scope). The maximal system resolution we achieved was 640×480 pixels, and the maximal frame rate was 25 fps. At maximal magnification, our field of view was 1.2 mm wide (or 533.3 pixels/mm, Supplementary Figure 2). The focus dial present mid unit becomes a zoom dial when the red zoom button on the unit is depressed. Our tracker has vertical slits which allow changes in magnification. Operating at maximal magnification and speed the videos obtained by the system are suitable for the study of various commonly measured physiological processes in the worm including digestion, defecation, egg laying, and pharyngeal pumping in single adult animals (Supplementary video 3). At the lower magnification range, we could film populations of worms with a visual field 10mm wide (80 pixels/mm). This resulted in adult worms measuring an average of 86 pixels in length. Tierpsy is the software package commonly used in professional settings and requires worms to measure a minimum of 100 pixels in length to obtain a full kinematic readout of animal postures. At intermediate magnifications our system achieves this benchmark, opening the door for using these movies for more in-depth analysis than what is available with the (easier to us) wrMTrck plugin for ImageJ.

##### Tracking

We filmed different *C. elegans* strains crawling and swimming. We next processed the avi movie files using the wrMTrck ImageJ plugin as described by the authors (Nussbaum-Krammer *et al.* 2015). To get an idea of how accurate the wrMTrck plugin was at tracking worms in our movies we multiplied the number of frames in each movie by the number of animals present in each movie. This allowed us to generate the maximum possible number of tracked events in each experiment. For example, a 60 second movie of 8 worms filmed at 25 fps would theoretically result in a maximum of 12,000 counted events (60 sec * 8 animals * 25 fps = 12,000). We divided the total number of events detected by wrMTrck by the theoretical maximal to generate a quotient of detection (QD) which ranges between 1 (when all animals are tracked the entire movie) to 0 (when no animal is tracked throughout the whole movie). Analyzing movies made with our tracker wrMTrck had a QD= 0.71+/−0.14SD). In our experience this quotient is similar to those obtained by professional software packages. The majority of the tracking error is the result of animals temporarily contacting the edge of the frame, momentarily exiting the field of view, or when animals collide and their outlines merge. Under these conditions animals are ignored by the tracking plugin (a common challenge for many machine vision algorithms). To illustrate the performance of the system we included supplementary videos 4-8 which show the successive steps of image processing and tracking for a group of swimming *unc-31(e928)* mutants. The tracking for this example had a QD of 0.84 resulting from occasional animal outlines merging. wrMTrck calculates many useful parameters such as body bends per second (frequency) and body lengths per second (normalized velocity) for each track that do not require an animal to be continuously tracked the duration of the video. In the example provided below we report swimming frequency and normalized velocity for the weighted average of the tracks obtained from each movie. One limitation of the wrMTrck plugin arises from the way the algorithm uses posture ratios to calculate body bend frequency. The lateral to longitudinal ratio used to detect body bends makes this parameter accurate only when analyzing swimming behavior when the posture of the worm permits the formation of a single body wave at the time. During crawling multiple waves travel posteriorly down the worm’s body and the algorithm consistently overestimates the number of body waves produced.

##### Cost benefit

Clearly investing in stronger and more durable materials will increase longevity and ease of use. We focused on producing a device that could be easily assembled, inexpensive, and that could last at least one semester of regular classroom use. Because of this, our system is not amenable to repeated changes of magnification settings. These changes are achievable using the vertical microscope slits that permit sliding the microscope up or down. We recommend having a couple of units dedicated to either high or low magnification if this is something the user anticipates needing regularly. Even with repeated changes in magnification we note that our builds made with thick corrugated cardboard lasted an entire semester in our undergraduate lab course. We also note additional limitations of our system compared to more expensive counterparts. More expensive (>US$1,500) systems like the one in use at the Haspel lab use infrared light and cameras to avoid potentially disrupting behavior. The cameras in such systems are high resolution and able to obtain high frame rates (>50fps), wider fields of view (>3cm), and high spatial resolution (>1900 pixels). These advantages allow users to film a hundred or more animals in a 5cm petri dish while still obtaining the high resolution needed by the Tierpsy worm spining algorithm. The larger field of view of these more expensive systems also allows the performance of behavioral assays (e.g. chemotaxis assays etc.). While our system is able to film worms at the resolutions required by Tierpsy, the smaller field of view necessitated by this magnification hinders combining this with behavioral assays such as chemotaxis, etc.

#### Sample Application

To illustrate the use of the worm tracking system we compared day one adult wild-type (N2) *C. elegans* worms to day 1 adult *unc-31(e928)* mutants. UNC-31 is expressed in synapses where it is predicted to have calcium binding activity. Loss of function mutations in this gene result in impaired in chemical synaptic transmission and severe locomotor defects (Speece et al., 2007). We set up our assays by placing eight animals inside a copper ring (1cm across) on an unseeded NGM plate. Copper repels worms and prevents animals from exiting the field of view during the assay (Simoneta and Golombek 2006). We next flooded the copper ring with 50 µl of liquid NGM to induce swimming. After two minutes of acclimation, we used the tracker to film worms for 60 seconds at 25 fps using our worm tracking system (Figure 1B). We repeated the experiment three times for each condition. Movies were exported as uncompressed AVI files (UYVY codec, Supplementary video 4), and uploaded to ImageJ. We used the “set scale” function in ImageJ to enter the appropriate magnification. We next used the scroll bar on the movie window to establish the limits of maximal vertical and lateral worm excursion. We then used the rectangular area of interest tool to select a region that was large enough to include the trajectory of all the animals during the entire movie and cropped the video to this smaller area in order to reduce file size (and exclude the copper ring or other visual artifacts) and converted the movie into 16bit (Supplementary video 5). Following wrMTrck instructions we generated a maximum intensity z projection picture of the movie, and then used the image calculator function to find the difference between the maximum projection and the movie. This effectively removed the background from the movie, and left only the moving (worms) objects (Supplementary video 6). In the final pre-processing step we adjusted the threshold of the movie image using the suggested maximum entropy setting and moving the dials to ensure that only the worms were selected. Care was taken by scrolling through the movie that the thresholding remained accurate throughout the movie. When applying the threshold, we selected the option “dark” background in order to produce a new movie with dark worms in a light background which is what wrMTrck requires (Supplementary video 7). At this point we were ready to apply the wrMTrck plugin. Before applying the plugin it is important to perform some initial measurements on the original movie file in order to establish the size of worms in pixels (as this is what the plugin uses in its measurements). Running the wrMTrck plugin we obtained a table with measurements and several other useful quality control features such as a movie with the tracked objects (Supplementary movie 8), and a bend track plot which allows for the empirical determination of the correct bend threshold to be used in body bend cycle determination.

We followed the processing described above for the N2 and *unc-31(e928)* strains and then focused on the body bends per second (i.e. swimming frequency), and body lengths per second (swimming velocity normalized to size) as these parameters are accurate descriptors of locomotor performance. The algorithm terminates a track whenever animals intersect or whenever they contact the edge of the visual field. Therefore there are more tracks than animals. Averaging the tracks would result in animals with fragmented trajectories being overrepresented. We therefore calculated a weighted average for each parameter of interest. To do this we took each track and multiplied the number of frames it lasted by the average measured reported for the parameter of interest (e.g. body bends per second). We then summed the products for all the tracks in the movie, and divided this by the total number of frames tracked (for all animals) in the movie. The resulting value is the average for the parameter in question, which assigns each contributing track a weight proportional to the fraction of the movie it lasted (i.e. shorter tracks having less weight than longer ones). We then plotted these values and used t-tests to compare the weighted averages of four assays of each strain for normalized swimming velocity and frequency. As previously shown (Speece et al., 2007), *unc-31(e928)* animals have severe locomotor impairments compared to wild types (Figure 1D and E respectively).

#### Concluding remarks

We described a worm tracking system that introduces an affordable, easy to construct and operate device intended for classroom use but capable of delivering results approaching those of more expensive professional systems. It is powerful and versatile enough to automatically measure common locomotor parameters, and to allow the study of physiological processes such as pharyngeal pumping, defecation, and egg laying which require high magnification and contrast. Although we did not test the system with other species, the system is likely useful for tracking other small organisms such as arthropod larvae, or even paramecia. We believe that the affordability and ease of use of the system will make it a useful addition to classrooms where physiological and behavioral consequences of gene and cellular activity are taught or investigated.

### Reagents

#### Organisms

We used 24 adult (day 1) wild-type (N2) *C. elegans* worms, and 24 *unc-31(e928)* adult (day 1) mutant worms (CB928) obtained from the Caenorhabditis Genetics Center (CGC), which is funded by the NIH Office of Research Infrastructure (P40 OD010440). Animals were raised under standard lab conditions (Brenner, 1974) at 20°C and fed on OP50 *E.coli* until used.

#### Statistics

We used SigmaPlot 11(Systat) t-tests to compare the swimming velocity and frequency for both N2 and *unc-31(e928)* worms using the weighted averages for four replicates with eight animals per assay.

#### Hardware

The following hardware was used to construct the worm tracking system:

A Windows computer may be required for filming, and it is required for analysis (not included in cost calculation).
USB microscope. US$20. Can be obtained from Amazon (ASIN: B08BFJZJ6N)
USB LED light. US$9. One needed (but comes in 8-pack) from Amazon (ASIN: B088KPZYS9)
USB LED dimmer. US$9. Suggested. Can be obtained from Amazon (ASIN: B01I17OHVQ)
Diffuser fabric. US$8. Enough for many setups. Can be obtained from Amazon (ASIN: B01JGY8U66)
Total build US$46 (US$31/ea if eight units are built)

#### Software

For use with the USB microscope there are many freely available software including Plugable Digital Viewer: https://plugable.com/drivers/microscope/. For this manuscript we used Windows onboard Camera software. There are differences in the level of control afforded by each software package. We recommend the user experiment with a couple free programs to determine which one suits their needs best.

To run the worm tracking software, it is necessary to first install ImageJ which is an open source Java image processing program found here: https://ImageJ.net/Fiji/Downloads We have used a few different versions of the program (ImageJ 1.53e and ImageJ 1.52p) without problems.

Worm tracking in ImageJ is accomplished by the wrMTrck plugin created by Jesper Pedersen available here: http://www.phage.dk/plugins/wrmtrck.html. A useful set of written and video tutorials are available online here: (http://www.phage.dk/plugins/download/wrMTrck.pdf and here: https://www.youtube.com/watch?v=V9M0s0E-0uI respectively.

## Acknowledgments

We thank the Caenorhabditis Genetics Center (CGC), which is funded by the NIH Office of Research Infrastructure (P40 OD010440), for providing strains.

## Extended Data

Supplementary Figure 1. Image. Supplementary Figure 1.pdf.

Supplementary Figure 2. Image. Supplementary Figure 2.jpg.

Supplementary Figure 3. Image. Supplementary Figure 3.tif.

Supplementary video 6. Audiovisual. Supplementary video 6.avi.

Supplementary video 5. Audiovisual. Supplementary video 5.avi.

Supplementary video 7. Audiovisual. Supplementary video 7.avi.

Supplementary video 8. Audiovisual. Supplementary video 8.avi.

Supplementary video 3. Audiovisual. Supplementary video 3.avi.

Supplementary video 4. Audiovisual. Supplementary video 4.avi.

Supplementary video 2. Audiovisual. Supplementary video 2.avi.

Supplementary video 1. Audiovisual. Supplementary video 1.avi.

Supplementary captions. Text. Supplementary materials.docx.

## Author Contributions

Nicholas Leonard: data curation, investigation, methodology. Andrés G. Vidal-Gadea: conceptualization, methodology, funding acquisition.

## Copyright

© 2021 by the authors. This is an open-access article distributed under the terms of the Creative Commons Attribution 4.0 International (CC BY 4.0) License, which permits unrestricted use, distribution, and reproduction in any medium, provided the original author and source are credited.

## Citation

Leonard, N; Vidal-Gadea, AG. (2021), Affordable *Caenorhabditis elegans* tracking system for classroom use. microPublication Biology.

