## Supplementary figures and images for "Affordable *Caenorhabditis elegans* tracking system for classroom use"

### Supplementary Figure 1

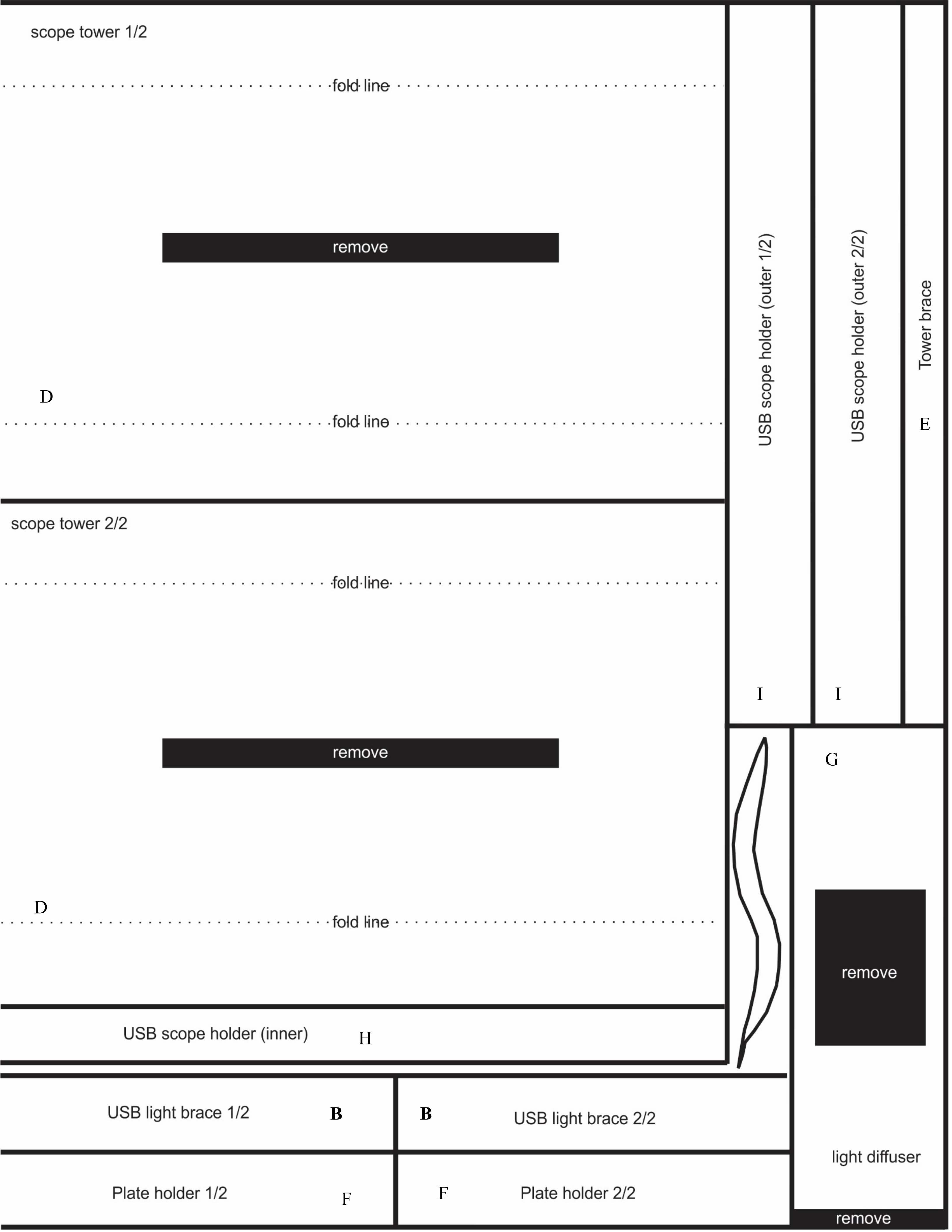

C

fold line

remove

fold line

remove

discard

A

fold line

fold line

### Supplementary Figure 2

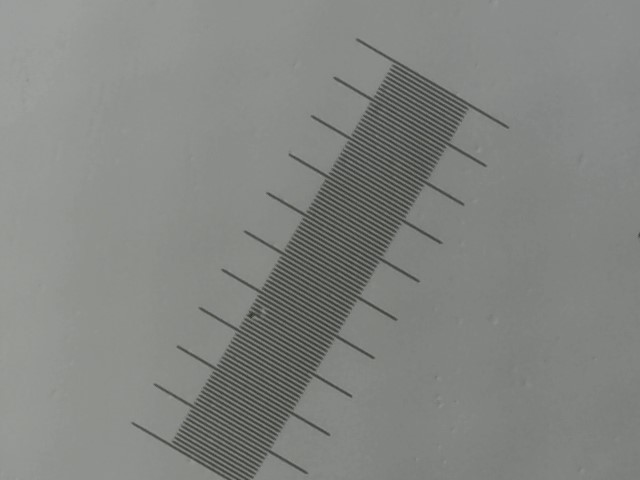

### Supplementary Figure 3

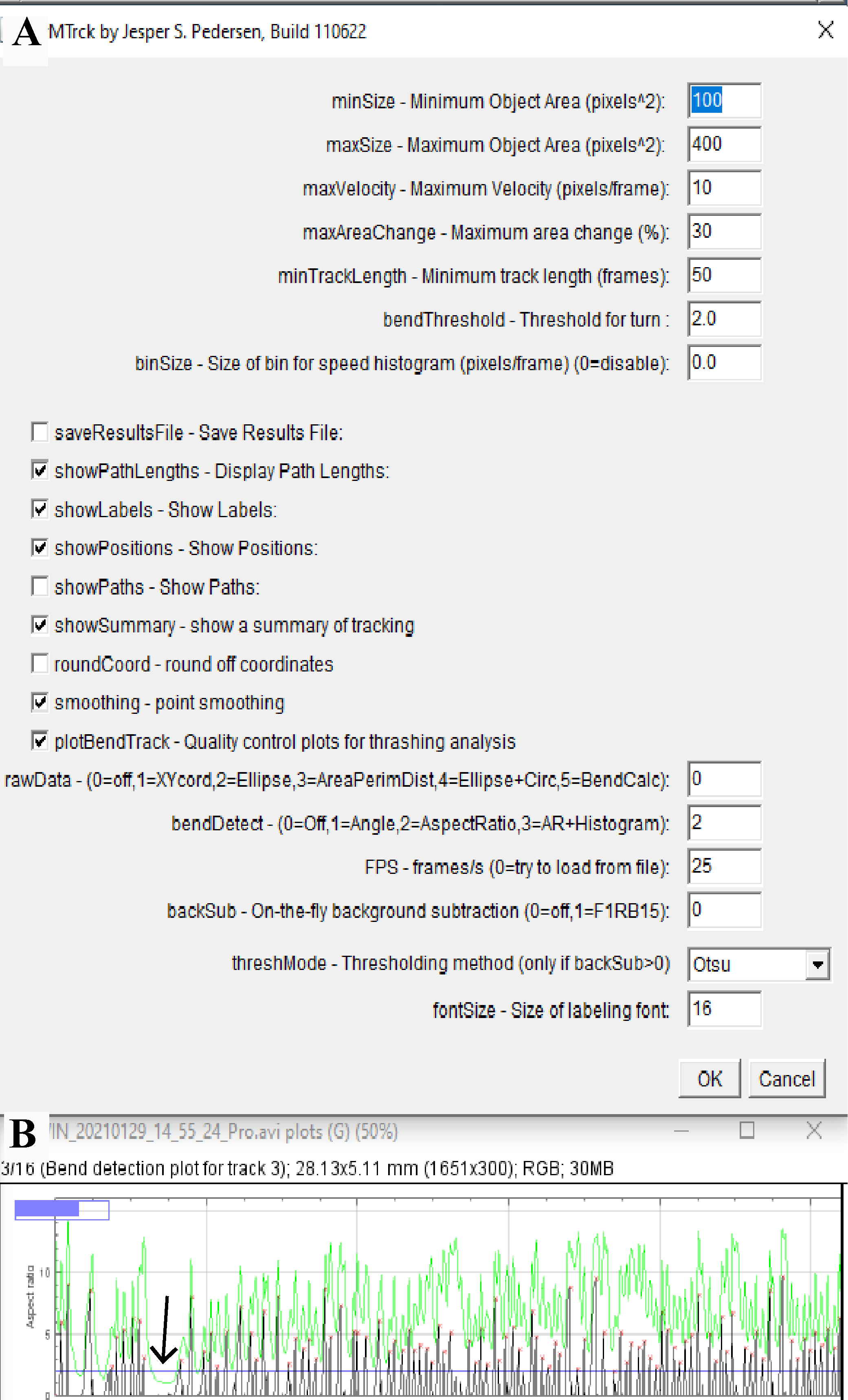
